# A systematic method for surveying data visualizations and a resulting genomic epidemiology visualization typology: GEViT

**DOI:** 10.1101/325290

**Authors:** Anamaria Crisan, Jennifer L. Gardy, Tamara Munzner

**Author notes:** Correspondence should be addressed to TM.

## Abstract

**Motivation:** Data visualization is an important tool for exploring and communicating findings from genomic and healthcare datasets. Yet, without a systematic way of organizing and describing the design space of data visualizations, researchers may not be aware of the breadth of possible visualization design choices or how to distinguish between good and bad options.

**Results:** We have developed a method that systematically surveys data visualizations using the analysis of both text and images. Our method supports the construction of a visualization design space that is explorable along two axes: *why* the visualization was created and *how* it was constructed. We applied our method to a corpus of scientific research articles from infectious disease genomic epidemiology and derived a Genomic Epidemiology Visualization Typology (GEViT) that describes how visualizations were created from a series of chart types, combinations, and enhancements. We have also implemented an online gallery that allows others to explore our resulting design space of visualizations. Our results have important implications for visualization design and for researchers intending to develop or use data visualization tools. Finally, the method that we introduce is extensible to constructing visualizations design spaces across other research areas.

**Availability:** Our browsable gallery is available at http://gevit.net and all project code can be found at https://github.com/amcrisan/gevitAnalysisRelease

## 1.0 Introduction

Genome sequencing is becoming an integral part of modern infectious disease diagnostics (Pankhurst *et al.*, 2016) and epidemiology (Faria *et al.*, 2016; Quick *et al.*, 2016). When genomic and/or phylogenetic data are combined with clinical and epidemiologic data routinely generated by public health laboratories and programs, the resulting analyses support a variety of public health professionals, including clinicians, epidemiologists, researchers, and policymakers, in their real-time decision-making around treatment, surveillance, and outbreak response. However, this new data-driven approach to public health also introduces interpretability challenges – it is difficult to succinctly and accurately represent such multivariate and high-dimensional data, particularly when many stakeholders do not routinely work with the genomic or phylogenetic data these analyses rely upon. These challenges arise not only late in an investigation, when attempting to communicate the results of an analysis, but also in the early phases of a project, such as initial data exploration and model-building (Grolemund and Wickham, 2014).

Data visualization is an important means to address interpretability challenges, and one which is increasingly being used in genomic epidemiology. Tools including nextstrain (Hadfield *et al.*, 2018) and Microreact (Argimón *et al.*, 2016) use developments in web technologies to produce sophisticated, interactive data visualizations that allow users to explore and interact with public health phylogenetic data in an epidemiological context. Other tools, such as treeviewer (Huerta-Cepas *et al.*, 2016), GenGIS (Parks *et al.*, 2013), or libraries such as PhyloCanvas, (http://phylocanvas.org/) also allow researchers varying degrees of freedom to generate visualizations blending phylogenetic trees with other metadata. As more and more visualization tools and libraries are being developed for genomic epidemiology, it is an appropriate moment at which to assess the type of visualizations being generated and used in public health genomic studies in order to inform the design of future visualization tools.

When analyzing existing data visualizations, the concept of a *visualization design space* becomes important. This design space is defined as the combinatorial space of data visualizations afforded by graphical marks (points, lines, and areas) that convey information through their aesthetic properties (position, color, size, shape, texture), which are also referred to as *channels* in the information visualization research literature (Munzner, 2014). There have been explicit attempts to describe visualization design spaces and share them via web galleries, such as SetVis (Alsallakh *et al.*, 2014), TreeVis (Schulz, 2011), Visualizing Health (https://www.vizhealth.org), and BioVis Explorer (Kerren *et al.*, 2017), but these were not created through a process as systematic as what we propose and thus do not serve to provide insight into current practice in a specific research community. Collections of visualizations also arise implicitly from search engine results, including Google, PubMed, or Semantic Scholar image searches, but these lack a systematic taxonomy and ontology describing the visualizations themselves. It is only through organizing the visualizations created by a research community with a design space that trends or common practices within that community become apparent and better practices can be suggested.

Here, we present a method for the systematic analysis of a visualization design space. By employing this structured approach to both generating and analyzing a suite of visualizations within the context of public health genomic epidemiology, we reveal current data visualization practices common to the field. We are able to identify those visualization designs that could be better supported through new software tools or improved to make them more effective, as well as areas of the design space that are currently underused. This methodological contribution can be applied to visualization design spaces in domains beyond public health genomic epidemiology; here we describe its application in a specific domain as an additional contribution. We present the Genomic Epidemiology Visualization Typology (GEViT), and we provide a web-based platform for exploring GEViT that researchers, bioinformaticians, and software developers can use to inform their own genomic epidemiology data visualization practice.

## 2.0. Materials & Methods

### 2.1. Developing a Method for the Systematic Analysis of Data Visualizations

Data visualizations are often challenging to analyze because, unlike images of real-world objects, visualizations in the scientific literature are abstractions devised by researchers to convey a combination of concepts. For example, phylogenetic trees display genomic data in an evolutionary context, and can be further enriched to show metadata about the sampled sequences and/or organisms and the underlying evolutionary processes. Visualizations vary across research contexts, and can be described using the nested model for visualization design and analysis (Munzner, 2009), which deconstructs a data visualization into four layers: the *why –* a research or domain problem that a data visualization supports; the *what –* the data that needs to be visualized and the specific tasks performed using the data and visualization, such as finding trends or communicating a specific finding; the *how –* the visual design and interactivity; and the algorithmic implementation of the visualization.

We have constructed a method for the systematic analysis of data visualizations that specifically articulates and then attempts to connect the visualization research problem (*why)* with the visualization design (*how)* – this goal is possible because we can meaningfully capture and label these elements of a data visualization through a systematic analysis based on image and textual analysis.

Our method consists of an initial literature analysis phase followed by a visualization analysis phase, resulting in a visualization design space in which images are classified according to their *why* and their *how*. The literature analysis phase (Figure 1) automatically analyzes text from a corpus of research articles to identify the topic of a data visualization – why it was created – as we assume that different topics are likely to yield different visualization designs. In the current instantiation of this method, we also use the literature analysis phase to perform a random stratified sampling of articles to select a reasonable subset of visualizations for the subsequent visualization analysis phase, which requires a human-curated inventory of each image. In this phase, we iteratively apply open and axial qualitative coding techniques to the set of images harvested from the sampled articles. The iterative qualitative coding phases (Charmaz, 2006) ultimately yield a set of hierarchical taxonomies that we collectively refer to as a *visualization typology* and that allows us to articulate *how* visualizations are created (Figure 1). Further detail around the methods employed during both phases are provided below and in Supplementary Figure S1.

**Figure 1.**
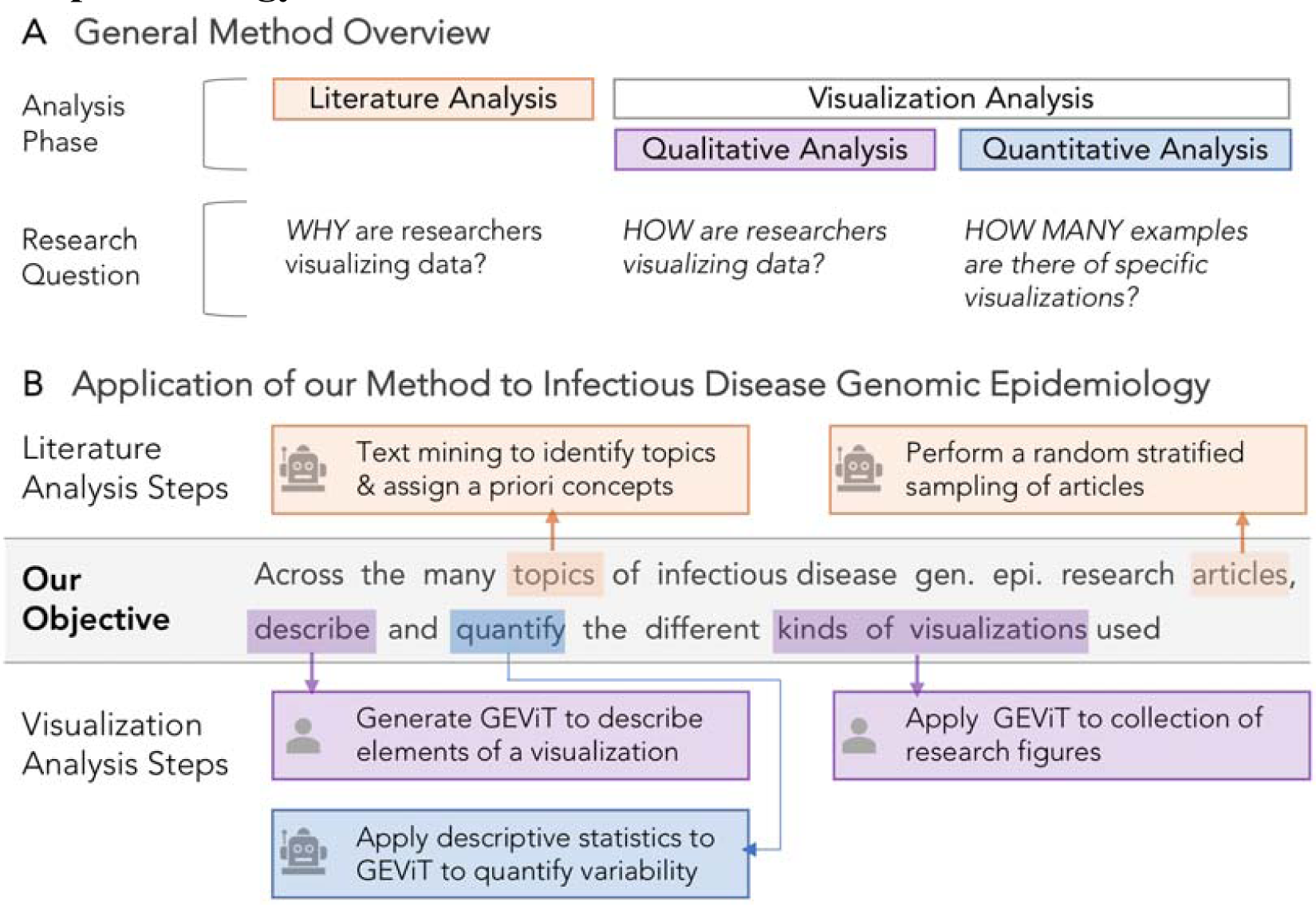
Method and Application Overview. **a)** Constructing and systematically analyzing a visualization design space requires analysis of both the literature and visualizations themselves, using qualitative and quantitative approaches. **b)** Automated steps, as indicated by the robot icon, are used in literature analysis to identify articles in genomic epidemiology and the topics those articles address. Manual steps, as indicated by the human icon, are used in the analysis of visualizations derived from those articles, followed by further quantification with automated statistical approaches. More details are also presented in Figure S1.

Our specific application of this method to articles and images from the infectious disease genomic epidemiology context resulted in the <u>G</u>enomic <u>E</u>pidemiology <u>Vi</u>sualization <u>T</u>ypology (GEViT) – a structured way of describing a collection of visualizations that together form a visualization design space. As a research community publishes new data visualizations, these can be annotated using the typology and added to the design space, and may even result in the addition of new terms to the typology if the image includes new elements of visual design.

### 2.2. A Systematic Analysis of Data Visualizations from the Infectious Disease Genomic Epidemiology Research Literature

All analysis notebooks and datasets are available online at:

https://github.com/amcrisan/gevitAnalysisRelease

#### 2.2.1. Literature Analysis

We developed an R package called Adjutant (Crisan *et al.* 2018b), described in detail elsewhere, to support our literature analysis.

##### Search Terms

We searched for articles related to infectious disease genomic epidemiology that were published within the past ten years. We used two queries, 1) *(genome AND (outbreak OR pandemic OR epidemic)) OR "genomic epidemiology"* and 2) *(genomic epidemiology OR molecular epidemiology) AND (bacteri* OR vir* OR pathogen) AND Genome,* combining their results and retaining only unique records for further analysis. We also manually included cancer genomics articles that were known to us to use phylogenetic trees in their analysis.

##### Data Preparation

The resulting document corpus included PubMed IDs, year of publication, authors, article titles, article abstract, and any associated Medical Subject Heading (MeSH) terms. Titles and abstracts were decomposed into single terms, stemmed, and filtered by Adjutant. We calculated the term frequency inverse document frequency (td-idf) metric for each term, and created a sparse Document Term Matrix (DTM) for further analysis. A separate dataset of bigram terms was also prepared and was used only to link articles to *a priori* concepts (see below).

##### Unsupervised Topic Clustering

We used the t-SNE and hdbscan algorithms to perform an unsupervised clustering using the DTM. We used the Barnes-Hut implementation of t-SNE (van der Maaten, 2014), which allows for some acceleration at the cost of accuracy, with the perplexity parameter set to 100; otherwise default parameters of the R package implementation were used (Krijthe, 2015). We then used hdbscan (Campello *et al.*, 2013) on the t-SNE co-ordinate to derive the topic clusters; we show in our earlier work on Adjutant (Crisan *et al.,* 2018b) that this order of operations yields relevant results. Clusters are sensitive to the minimum number of cluster points (minPts) parameter supplied to the hdbscan, thus we tried different minPts values (50, 75, 100, 125, 150, 250, 500, 1000), observing how the cluster compositions changed. We observed that some articles never held membership in any cluster irrespective of the parameter settings and labelled those as “never clustered”, in contrast to articles that were simply not clustered with our specific final parameter settings that are labeled as “currently unclustered”. The final set of clusters combined results from the minPts 75 and 150 analyses. Each cluster is assigned a topic by using the two most frequent terms within the cluster. Following topic clustering, we validated our clusters using an external list of human pathogens (Table S1), assessing the correspondence between pathogen names and cluster topics.

##### Linking To *A Priori* Concepts

Before conducting the unsupervised clustering, we discussed what results we might expect given our knowledge of research activities in the public health genomic epidemiology community. This initial discussion produced a set of 23 *a priori* concepts that we categorized into three groups: genomic concepts, including drug resistance, genome, genotype, molecular biology, pathogen characterization, phylogeny, and population diversity; epidemiology concepts, including clusters, disease reservoirs, geography, outbreaks (at international, community, and hospital levels), surveillance, transmission, vaccine, and vectors, and medical concepts (clinical, cancer, diagnosis, outcome, and treatment).

Following the clustering, we identified bigrams that occurred in at least ten articles within a pathogen topic cluster and between at least 10% of the other pathogen topic clusters, and manually assigned those bigrams to an *a priori* concept (Table S2) – for example, the bigram “vancomycin resistance” was assigned to the *a priori* concept of “drug resistance”. Assignments were validated by internal discussion among the research team, including a genomic epidemiology expert.

##### Document Sampling

To produce a manageable, diverse, and systematically derived dataset for the human-curated visualization analysis step, we performed random stratified sampling on our document corpus, sampling one document for each *a priori* concept within each of the automatically derived topic clusters. Each sampled article was examined and either considered acceptable for further analysis or rejected. Most articles were rejected because they did not contain any figures; other reasons for rejection included: full text article not accessible; article not in English; article was about a laboratory or bioinformatics technique and not an epidemiological scenario; no human data; or the article was a review rather than original research. For each rejected article, we resampled two additional articles, choosing one for further analysis. Based upon the analysis of the first round of sampling, the second round only sampled articles from 2011 onwards to increase the chance of sampling articles containing figures, and also attempted to sample underrepresented *a priori* concepts from the first round. Table S3 contains a list of all the articles, which round they were sampled in, whether they were included or rejected, and the reason for rejection.

##### Figure and Table Extraction

To properly capture the figures and their captions, we manually extracted them from PDFs of the sampled articles. Images were only excluded if they were CONSORT diagrams, flow diagrams, or illustrations without underlying data. We also included a small number of “missed opportunity” tables – stand-alone tables that we felt could have been visualized, most frequently matrices of numbers or large tables of patient metadata where each row consisted of a patient.

#### 2.2.2. Visualization Analysis

Extracted figures and tables were analyzed using iterative open and axial qualitative coding techniques. Originally derived from the use of Grounded Theory in sociology, psychology, and anthropology (Charmaz, 2006), qualitative coding methods are now being used in human-computer interaction (Jacko, 2012) and information visualization research (Carpendale, 2008). Qualitative coding involves iteratively examining data and assigning it to some category. The categories themselves are refined and can take on hierarchical relationships through different cycles of the coding process (see Supplemental Methods), and were informed here by concepts from visualization theory and terminology (Munzner, 2014).

Here, we analyzed whole figures separately – we did not decompose multi-part figures in order to understand the potential interplay between panels within a figure. We began by creating a taxonomic code describing the types of charts present in different figures. We next examined how different types of images were combined to show different aspects of the data and thus created a chart combination taxonomy. Finally, we created a taxonomy that captured how basic chart types were enhanced to encode additional information. We refer to the collection of taxonomic code sets for chart types, combinations, and enhancements that were derived from this document corpus of genomic epidemiology research articles as GEViT. We conducted three rounds of qualitative coding, in which we reviewed figures and made additions or changes to GEViT; by the third round of coding, there were too few additional modifications to warrant a subsequent round.

#### 2.2.3. Creating an Explorable Visualization Design Space

We used the results of the literature and the visualization analysis phases to produce an explorable visualization design space, which is freely available at http://gevit.net. The images presented gallery are used under Fair Use copyright terms and we provide links back to the original source publications.

## 3.0. Results

### 3.1. Literature Analysis

#### 3.1.0. Literature mining identified article clusters according to disease pathogen

We assembled a document corpus of 17,974 articles pertaining to infectious disease genomic epidemiology research published in the past 10 years (Figure 2). Using article titles and abstracts we derived topic clusters in an unsupervised manner, and classified articles as either belonging to a named topic cluster, not belonging to a cluster under current parameter settings, or never being clustered under any parameter settings (Figure 3a). Articles that never formed part of a cluster were removed from further analysis, leaving 15,315 documents of which 11,416 (75% of the initial document corpus) formed 32 topic clusters (Figure 3b). Clusters were assigned topics via the top two most frequent terms within the cluster, revealing that infectious disease genomic epidemiology literature is primarily structured around pathogens. We validated our results by comparing our automatically derived cluster naming to the distribution of pathogen terms from an external list (Table S1, Figure 3c), and found there to be a strong correspondence between the automatically derived cluster topics and the propensity for pathogen terms to appear within clusters of the same name (for example, the term “*influenza virus*” occurs primarily within the “influenza-viru” cluster). Some notable exceptions are *Escherichia coli, Helicobacter pylori*, and *Human Immunodeficiency Virus* – in addition to having their own defined cluster, these terms also appeared in other clusters, suggesting co-infections or another phenomenon. We also found that clusters with more generic names (for example “viru-sequenc”, or “geno-sequenc”) contain pathogens that likely had too few articles to form their clusters, possibly reflecting recently emerged pathogens (i.e., Zika, Ebola) with a less extensive research history. We filtered the corpus by limiting to pathogens with 40 or more articles, resulting in 6,350 articles within 35 pathogen clusters, then further simplified to 18 clusters: a final set of 17 pathogen clusters that had 100 or more documents and one “other” cluster (Table S4).

**Figure 2.**
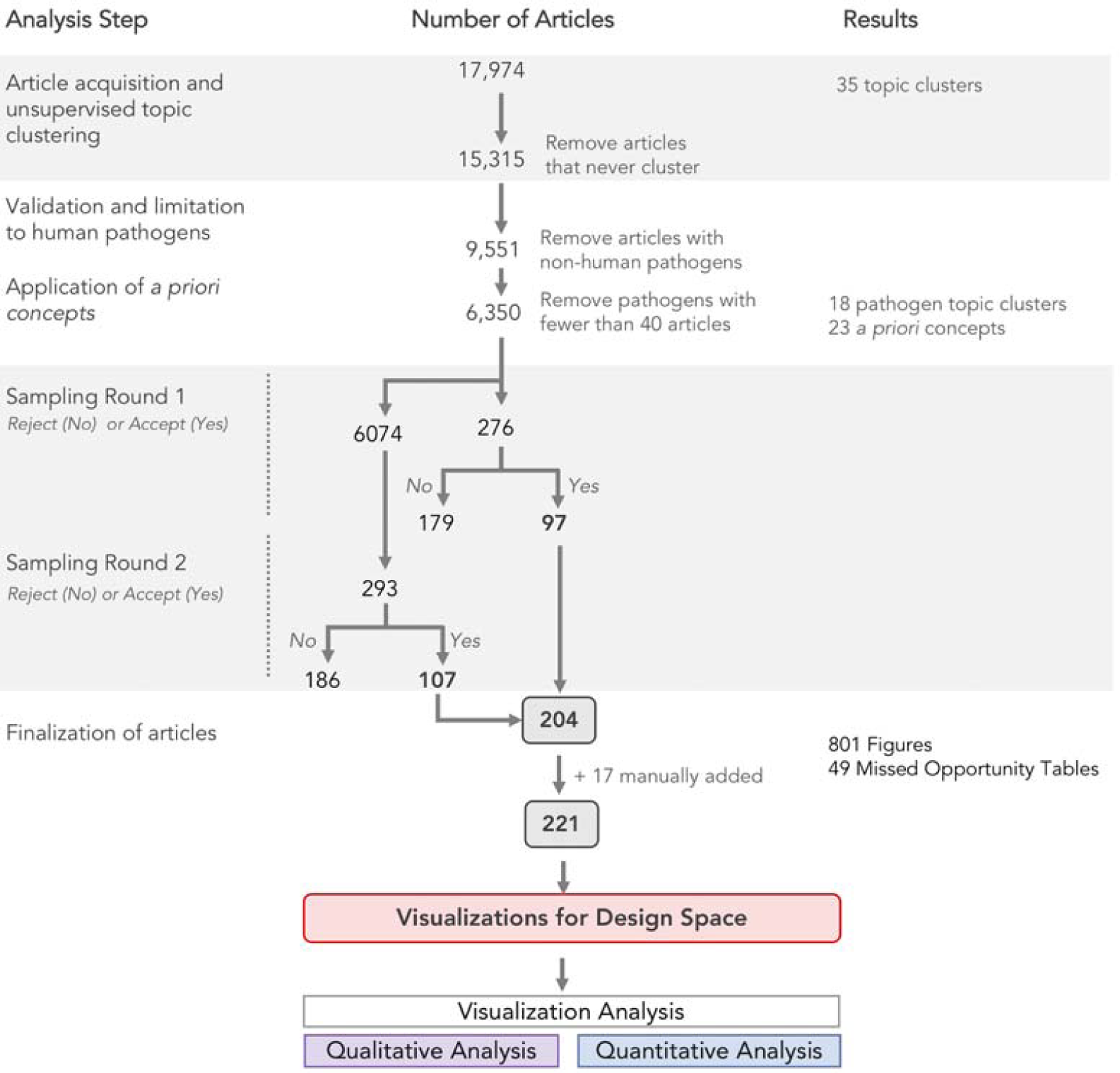
Summary of literature analysis steps and document sampling.

**Figure 3.**
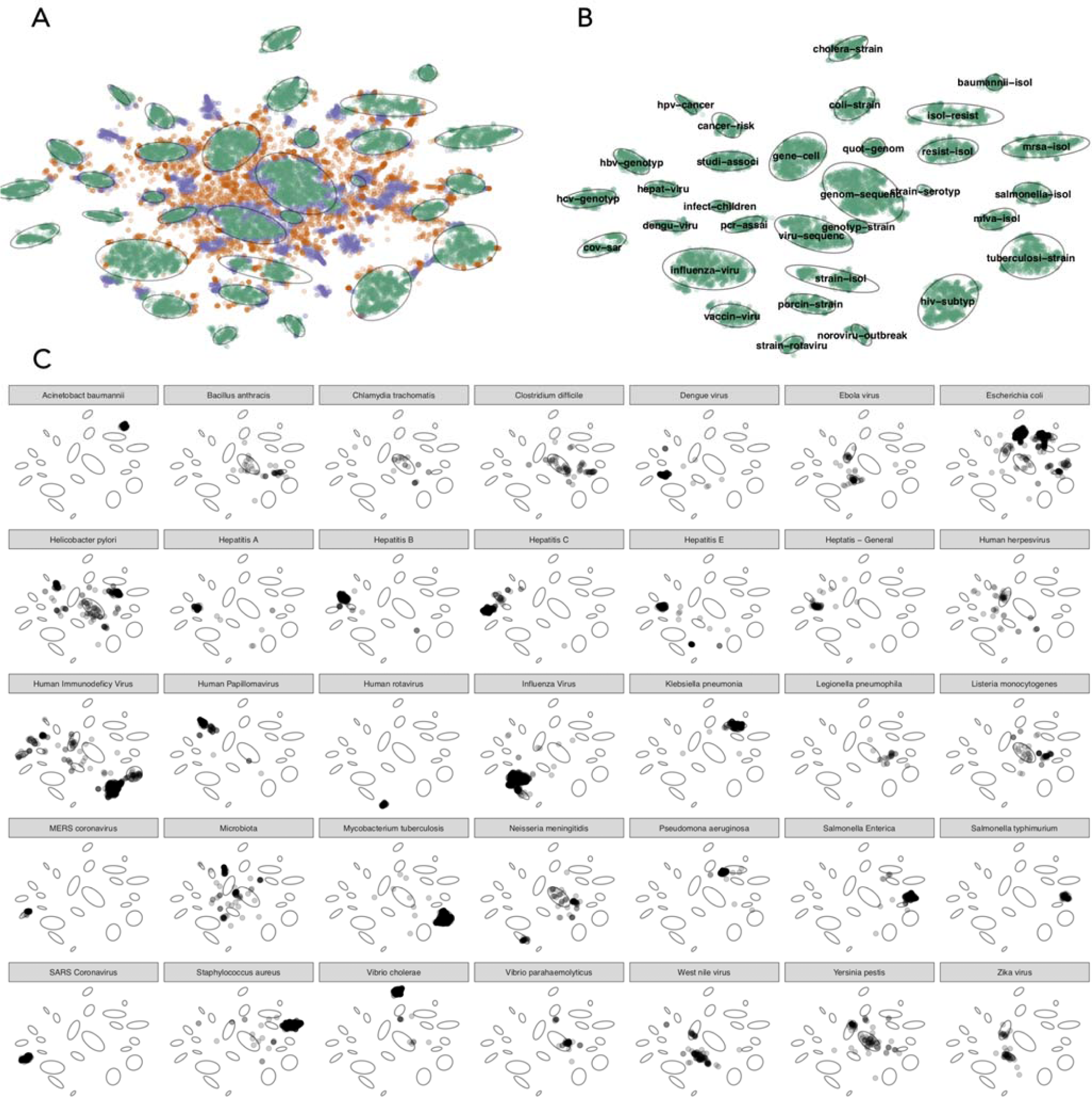
Summary of literature analysis results. **a)** Documents were classified according to whether they were part of a cluster (green), unclustered under current parameter settings (purple), or never clustered (orange). The 32 cluster boundaries were automatically determined and are shown as light grey ovals. **b)** Clustered documents and their topics, which are automatically assigned based upon top two terms with the cluster. **c)** Verification of cluster topics against an external list of pathogens. The small multiples show the distribution across the clusters of the pathogen named in the panel header, for the 35 pathogens with 40 or more matching documents.

#### 3.1.1. Clusters were manually mapped to *a priori* concepts

The findings from the literature mining were at odds with our own *a priori* assumptions that articles would cluster according to more general, pathogen-agnostic concepts, such as drug resistance, surveillance, and outbreak investigation. In order to allow researchers to investigate the connections between these familiar *a priori* concepts and the literature-derived clusters, we linked them together manually. We mapped a total of 23 *a priori* concepts to 404 bigrams. We found that *a priori* concepts did not occur uniformly across pathogen clusters (Figure S2A) and a variable number of bigrams mapped to individual *a priori* concepts, with 143 bigrams mapped to “drug resistance” and only one bigram mapped to “disease reservoirs” (Figure S2B).

#### 3.1.2. Document sampling was stratified according to pathogen and *a priori* concepts

We then performed two rounds of stratified sampling using pathogens and *a priori* concepts as strata. The sampling resulted in 204 unique articles, to which we manually added 17 additional articles that we deemed contained interesting data visualizations mainly from cancer research – these are clearly tagged in our analysis – for a total of 221 articles (Table S1) from which we extracted a total of 770 figures, including a small number (45) of ‘missed opportunity’ tables.

### 3.2. Visualization Analysis

#### 3.2.0. Developing GEViT – A Genomic Epidemiology Visualization Typology

Using the analysis set of figures from the sample documents, we used iterative open and axial coding techniques to devise a systematic way to describe how data visualizations are constructed (see Supplemental Methods). We began by classifying the types of charts in figures, then classified how charts were combined, and then classified how charts were enhanced. We found that these three descriptive axes allowed us to sufficiently describe all visualizations in our dataset of figures. For each of these descriptive axes, we also derived a hierarchical taxonomy. Collectively, we refer to this result of the descriptive axes and their associated taxonomies as GEViT (<u>G</u>enomic <u>E</u>pidemiology <u>Vi</u>sualization <u>T</u>ypology). Below, we describe each of GEViT’s descriptive axes and interleave descriptive statistics to show the distribution of taxonomic codes across these axes to provide an overview of the variance in the resulting visualization design space.

#### 3.2.1. Chart Types in GEViT

We identified eight classes of chart types that form the basis of the data visualizations in our dataset (Figure 4): Common Statistical; Color (statistical charts that intrinsically depend on hue or brightness to convey data); Relational; Temporal; Spatial; Tree; and Genomic. We compiled a taxonomy of common chart names to classify specific instances of chart types within each class. When applicable, we also defined special cases of a specific chart – for example, epidemic curves are a special case of bar chart. We also defined one ‘Other’ category, which included entities that accompanied data visualizations but were not themselves data visualizations, such as tables and images, and miscellaneous visualizations that did not fit elsewhere. In total, we observed 23 distinct chart types plus one miscellaneous category and found that the most commonly occurring chart types within data visualizations included Phylogenetic Trees (17.7% of all data visualizations, although some type of tree was present in 23.7% of all visualizations), followed by Tables (9.7%), Bar Charts (8.9%), Genomic Maps (6.9%), Line Charts (6.8%), and Images (5.7%, typically a Gel Image of Pulsed Field Gel Electrophoresis) (Supplemental Figure S3). The frequency of tables, either alone or in combination with another chart type, is a notable finding, indicating missed opportunities for visualization. Our findings also suggest that only a small portion of the available design space is typically used.

**Figure 4.**
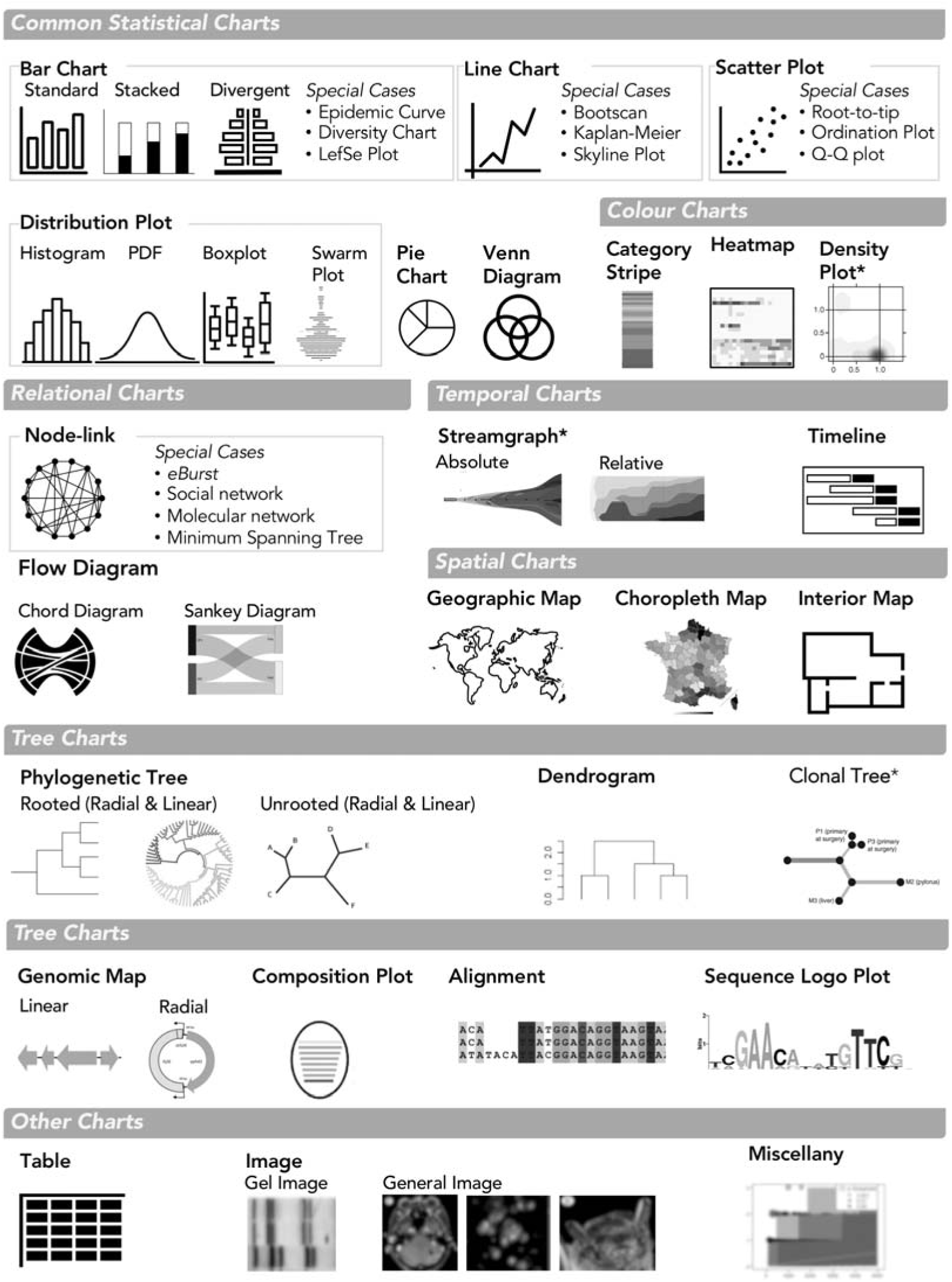
Chart Types in GEViT. We used common names for chart types and separated them into eight main classes and also one ‘Other’ class. Special cases of chart types were defined only when there were multiple instances of the same specific chart across our dataset. Chart types with an asterisk mark (*) indicate that they were included in the analysis through manually added articles.

#### 3.2.2. Chart Combinations in GEViT

The majority of figures were composed of a single chart type (40.1%), but we observed distinct and common patterns of combining chart types to create more complex, and often linked, multi-part figures (Figure 5). Composite charts (20.3%) contained multiple chart types that were spatially aligned – for example, a heatmap and dendrogram are spatially aligned to jointly convey clustering information. A tree and heatmap can also be visualized independently of each other, but their combined value is evidently relevant for many researchers. Small Multiples (17.3%) showed different aspects of the data through multiple instances of the same chart type. Many Types Linked combinations (13.5%) used multiple different chart types that were visually linked – for example, using a common color to denote some property of the data across the different charts, but not spatially aligned (in contrast to Composite charts). Finally, Many Types General combinations (8.8%) comprise multiple chart types, but where there is no spatial or visual link between charts – these likely were combined into a single figure due to manuscript space restrictions. It was not always straightforward to distinguish between some instances of Many Types Linked and Many Types General, and in such cases, we resolved the ambiguity in favor of the latter classification. We also observed instances of Complex Combinations (11.9%), in which visualizations used two of the previously described chart combinations. Phylogenetic Trees were the chart type mostly commonly combined with other chart types.

**Figure 5.**
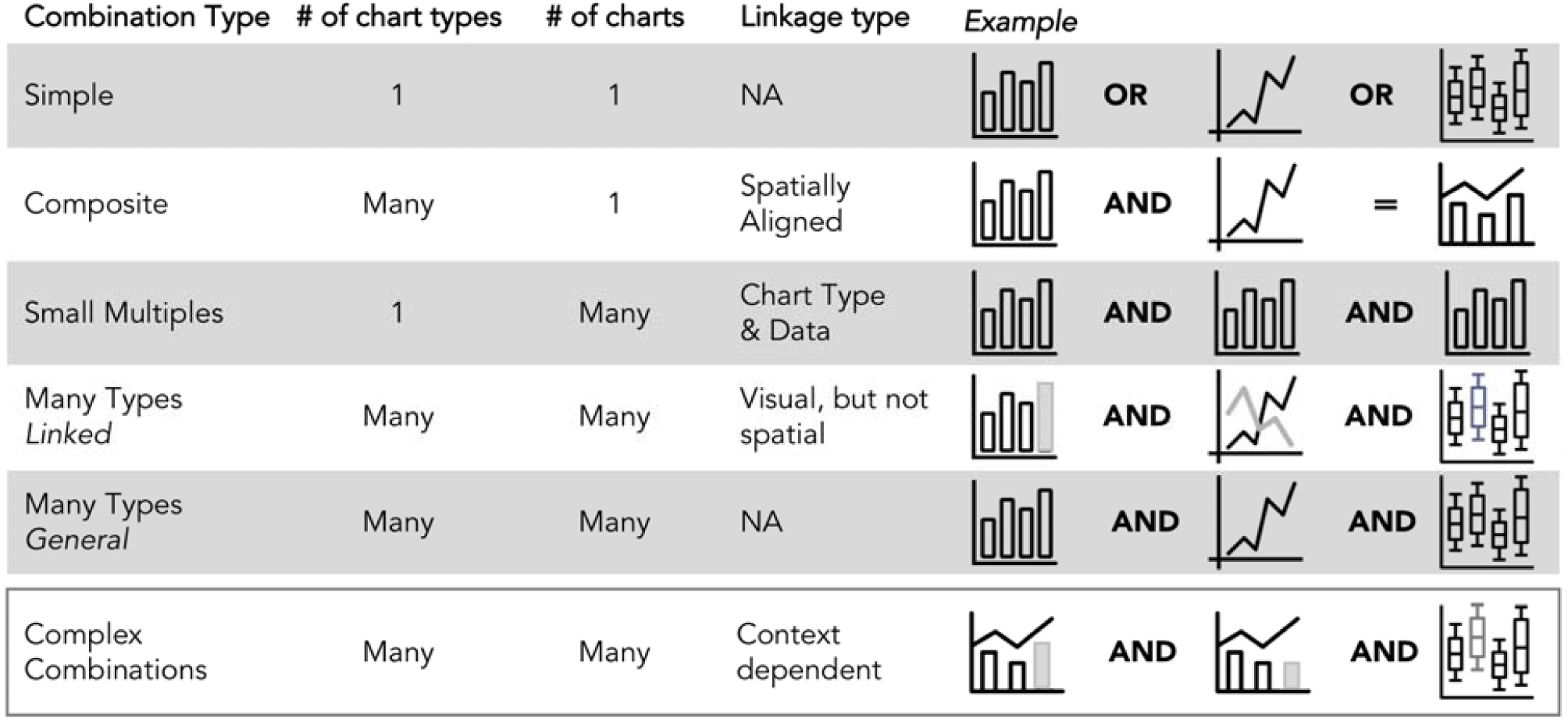
Chart Combinations in GEViT. The six combination types differ based on the number of chart types, the number of charts, and the approach to linking them together. Complex combinations are an amalgamation of the above five chart types – for example, a composite visualization that is represented as a small multiple and also linked another chart type.

#### 3.2.3. Chart Enhancements in GEViT

Lastly, we noted that standard chart types were often enhanced to add metadata through the addition or changing of graphical marks – the basic graphical element corresponding to a data record (*e.g.* a patient), or derived data value (*e.g.* the total number of patients). Graphical marks are points, lines, areas, and text, which are endowed with the aesthetic properties of size, shape, color, and texture that can be modified to encode data (Figure 6a). For example, a phylogenetic tree encodes evolutionary relationships inferred from nucleic acid or protein data as lines of some calculated length (Figure 6b). These lines are often black; however, they can be re-encoded to incorporate data from some additional source – for example, coloring lines according to geographic regions. It is also possible to add marks to the base chart type – for example, adding colored point marks to a tree’s leaf positions (Figure 6b), or adding linear brackets and text to delineate or otherwise annotate groups. We did not consider axis text, titles, or data labels to be added marks, subsuming them as constituent parts of the base chart type.

**Figure 6.**
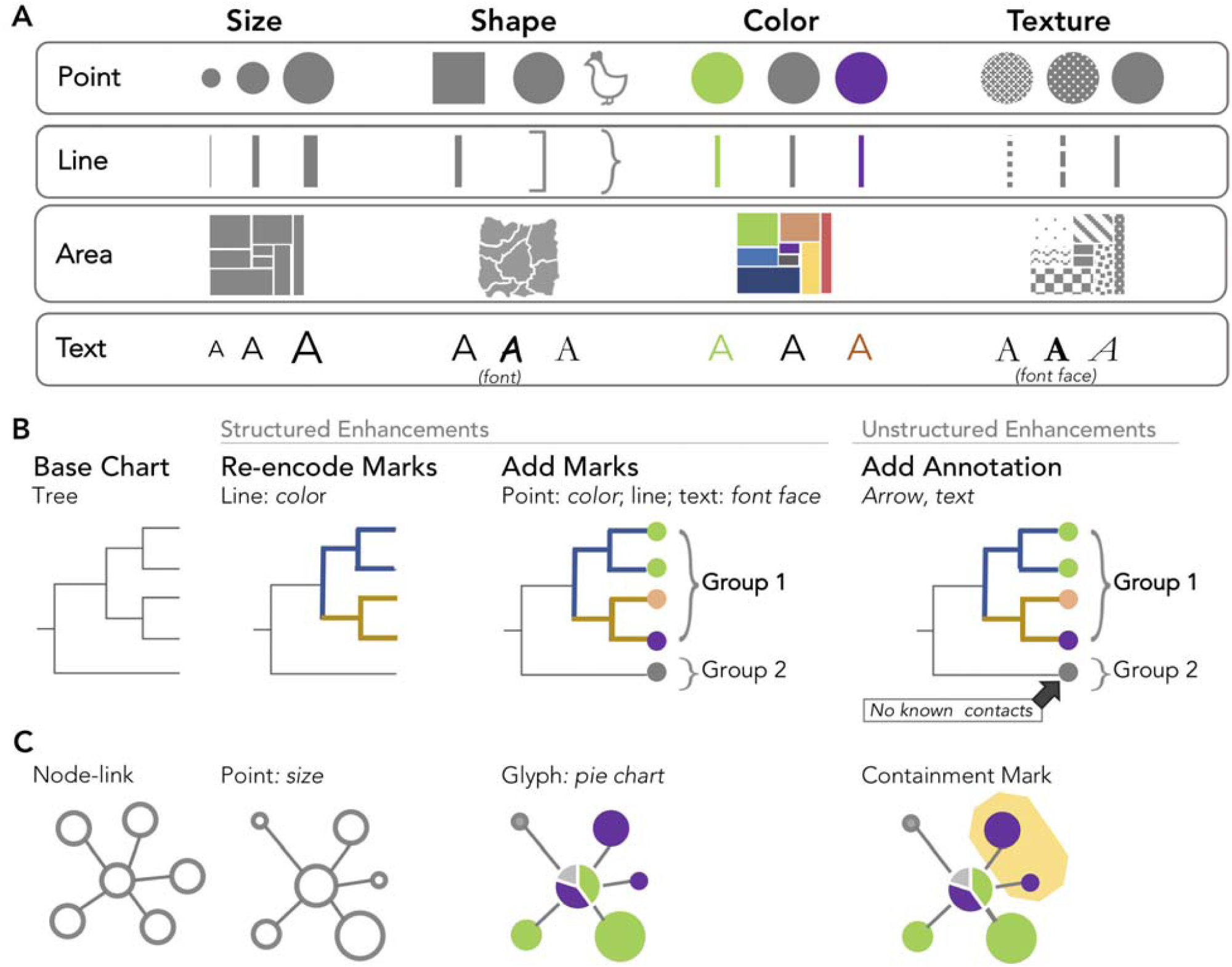
Chart Enhancements in GEViT. **a)** Our characterization of marks and their associated aesthetic properties is based on longstanding conventions in the visualization literature (Munzner, 2014; Meirelles, 2013) with roots in Bertin’s *Semiology of Graphics* (Bertin, 1983). Illustrative examples are shown for **b)** a tree and **c)** node-link chart types.

It is also possible to add more complex types of marks. Connection marks are a specific instance of line marks that connect two other marks. Containment marks are a specific instance of area marks that enclose other marks. Finally, a glyph is a complex mark that could itself be a type of chart, but that is smaller than the base chart type and embedded within it (in contrast, we define that composite chart types have the same frame size and one chart is not embedded within the other). The only glyph we identified within our dataset was a pie chart, which was often added to geographic maps or node-link graphs (Figure 6b) to communication proportions within the data.

We differentiate between the instances when chart enhancements are added consistently, or just as one-off marks. When the addition or re-encoding of marks is applied consistently to the base chart type – for example, re-encoding all or many lines in a tree or adding points to all or many leaf nodes – we defined these as structured enhancements. Adding one-off marks, even if they are driven by the data or the addition of some arbitrary ink, was considered to be an annotation and defined as an unstructured enhancement. It was not always easy to differentiate between structured and unstructured enhancements, and in such instances, we resolved ambiguities by choosing structured enhancement when analyzing figures.

In our dataset, we observed that most base chart types were enhanced (83.8% of all chart types), typically through the addition of lines, points, or text (59.6%), while re-encoding of marks was less common (45.6%). The use of text as a graphical mark with aesthetic properties that can be manipulated to convey information was common in our dataset, either by adding text marks to a base chart type, or re-encoding of text labels by manipulating the font face. The text itself ranged from the simple case of a single letter or number, to a full word, to a complex concatenated string of metadata such as specimen ID, location, and year. Annotations were also less common (33.6%), and were most commonly an arrow to text or a containment mark that highlighted only a single group.

#### 3.2.1 The GEViT Gallery – an Interactive Exploration of the Visualization Design Space

We created a browsable gallery, available at https://gevit.net, that allows others to explore the genomic epidemiology visualization design space we created, examine the results of the literature analysis, and browse our GEViT taxonomic code sets. Visitors to the GEViT gallery can browse visualizations across different pathogen types, contextual tags derived from *a priori* concepts, and data or arbitrary terms found in the figure captions. Clicking on an individual figure within the gallery reveals its construction via GEViT. Users are also able to browse visualizations based upon GEViT’s taxonomic codes to see the myriad applications for certain chart types, combinations, and enhancements. In our analysis of the data visualizations we also identified examples of good and bad visualization design practice.

## 4.0 Discussion

Data visualizations are important outputs of many scientific investigations and, when viewed collectively, merit close study to reveal common, good or bad, or missed practices in visualization design. Here, we describe a method for systematically studying data visualizations from the scientific literature, using both text and image data to articulate both *why* a visualization was created and *how* it was created from various chart types, combinations, and enhancements. We applied this method to a literature corpus from the domain of infectious disease genomic epidemiology, resulting in the <u>G</u>enomic <u>E</u>pidemiology <u>Vi</u>sualization <u>T</u>ypology (GEViT), and we created a web-based GEViT gallery, allowing users to browse the visualization design space generated through the application of our methodology and to see how elements of our typology can be used to describe a given image.

The typology aspect of our work is similar in spirit to the Grammar of Graphics proposed by Wilkinson (Wilkinson, 2010) and modified and instantiated by Wickham within ggplot2 (Wickham, 2010). While that prior work focuses on low-level details of chart implementation ours uses a higher level of abstraction, using whole chart types as a basis. This higher level of abstraction is more appropriate for exploring and describing the design space used by a community. Our work also differs from existing ontology-based efforts to describe a research domain, such as GenEpiO (Griffiths *et al.*, 2017). GEViT might be considered as the data visualization equivalent to the structured vocabulary that an ontology provides; however, it does not describe the relationships among entities as an ontology would. However, with future work, incorporating visual typologies into ontologies like GenEpiO is possible.

The present study used data visualizations from articles within the published research literature and did not include visualizations intended for public consumption, not published in peer-reviewed journals. This choice was pragmatic. First, it bounded the search space for our analysis. Academic articles are often accessible through specialized literature repositories, whereas including public-facing visualizations would have required extensive web scraping. Furthermore, research articles are relatively structured, making them an ideal substrate for topic modelling. Limiting our analysis to peer-reviewed scientific literature also bounded the content of the sampled images. There is shared technical knowledge within a research community, meaning that most users can interpret a visualization without additional assistance, whereas visualizations designed to communicate a concept to a more general audience often incorporate additional explanation or background information (although many of these more general, public-facing visualizations likely begin as images created in the academic research context). The typology we developed is extensible to more unstructured, non-academic data visualizations used for public communication, and it would be interesting to compare such a design space with the one we present here. An important limitation imposed by our literature search strategy is that we only included the final data visualization used to communicate some research finding – we do not have access to those data visualizations that researchers created during their internal data analysis process. Our own experiences in public heath genomics research and developing data visualizations to share our research findings suggest that the visualizations used during an analysis and those used to communicate final results do not substantially differ, but confirmation of this conjecture would be a good subject for future work.

Another limitation of our current method is the requirement for a human to manually carry out aspects of the visualization analysis. Although this process was time consuming, this inclusion of the human in the loop was crucial to understand what aspects of each visualization were necessary to delineate a design space; current machine learning methods are not capable of generating such a result. Developing a semi-automatic method that combines some automatically created decomposition of visualizations with human judgement as part of the analysis loop through future work would accelerate the process of refreshing and maintaining the GEViT resource.

### Implications of our findings for visualization design

By creating a visualization design space, we not only capture current common practices in our research domain, but we also reveal gaps and areas that require additional attention. While we found some instances of bespoke, effective, and aesthetically appealing data visualizations, the systematic nature of our method to exploring visualization choices reveals that across pathogens and *a priori* concepts, visualization design choices are quite homogenous, and the quality of visualizations varies substantially. We had expected greater variability, given than different pathogens have different transmission routes, are responsive to different interventions, and exist in different environmental, zoonotic, and human contexts. However, phylogenetic trees are the dominant visualization choice, often with additional contextual data included as tree labels or as accompanying tables. This dominance may impact effective knowledge translation in the genomic epidemiology domain, as the interpretability and utility of trees is unclear among public health decision-makers who have limited experience with genomic data (Crisan *et al.*, 2018b).

Although our finding of design homogeneity is not surprising, it also underscores how a lack of awareness of design alternatives leads to ineffective data visualizations. For example, geographic data is often encoded as text rather than an alternative mark or an explicit visual representation. The pervasive use of text in genomic epidemiology visualizations stands in contrast to recommendations from the information visualization research literature, where the use of text as a mark type is discouraged. Reading text requires more working memory and thus imposes a high cognitive load, whereas the goal of most visualizations is to reduce cognitive load by leveraging human perceptual systems to interpret information through the encoding of data as marks and aesthetic properties (Munzner, 2014). Our finding that text was often used as a mark and was endowed with aesthetic properties like color and variable font faces and sizes suggests that researchers are aware of the power of visual channels, but not necessarily the choice of an effective mark. We also note that many visualizations tended to show all of the data, rather than exploring alternatives that visually summarize data at multiple levels of detail.

Our work highlights opportunities for further work on areas where the genomic epidemiology research community could be better supported in designing data visualizations. Bioinformaticians and software developers can use GEViT to evaluate whether the tools they are creating afford the visual expressivity that infectious disease researchers need to communicate their research findings. Phylogenetic trees are evidently important, but there is a need for better tools that allow researchers to explore alternative visualizations and to more effectively encode tabular metadata onto trees and other visualizations. Although our study did not reveal how researchers create their data visualizations, our experience in the genomic epidemiology research community suggests that many chart- or tree-generating packages, some in R, are often used in conjunction with Power Point or Adobe Illustrator to compose complex visualizations that include chart combinations and enhancements. Software tools or libraries that support more expressive generation of visualizations can lower the barrier to generating data visualizations, reduce the overreliance on text, expand the use of combinatorial charts, and contribute to more reproducible research by creating more informative visualizations in which data is not obscured.

We also suggest that our findings might inspire developers to create alternatives to existing common design choices, and that our gallery of visualizations gives such developers a resource with which they can empirically test their new visualization design against existing choices. This empirical approach to testing a new visualization will help move the community further away from the *ad hoc* approach to visualization development, where design choices are heavily biased by individual preferences. As more work is done to explore and test new visualization designs, GEViT will incorporate these designs, potentially resulting in the addition of new typological terms. It will also be interesting to explore how GEViT might be used to suggest visualizations to researchers, as is currently done with common statistical charts in tools like Tableau’s “Show Me” feature (Mackinlay *et al.*, 2007), Google Sheets’ “Chart Suggestions”, or in novel systems like Draco (Moritz, 2018).

### Implications of our findings for the genomic epidemiology community

Data visualization can be an important tool for translating scientific results to a group of experts working in a common domain but with varying backgrounds. This situation is often the case in public health genomic epidemiology, where microbiologists, computational biologists, clinicians, epidemiologists, healthcare administrators, and others often come together around a specific issue. By making individuals aware of data visualization conventions used by the community through the GEViT gallery, we hope to assist researchers who struggle to visually communicate their research findings by providing both inspiration and a framework for reasoning about data visualizations that will assist as they develop their own data visualization practice. We have tagged examples in the GEViT gallery with “good” and “missed opportunities” to provide some guidance, but these labels are assigned by our subjective reasoning as data visualization experts and have not been empirically validated. Future work in this area might include quantitative evaluation of the efficacy of particular visualizations, and ultimately more sophisticated guidance around visualization design and analysis in the public health context.

## 5.0 Conclusions

Through a systematic method, we have delineated the visualization design space used in infectious disease genomic epidemiology. We provide both a concrete terminology for describing data visualizations and a gallery of visual inspiration, the combination of which we hope will provide guidance to visualization tool developers and to researchers looking to create their own visualizations. Mostly importantly, our work demonstrates that is possible to think systematically and rigorously about data visualizations and that there exist open, complex, and interesting problems in visualizations design and analysis, where the potential impacts on research domains such as public heath are profound.

## Acknowledgements

The authors appreciate discussions with the InfoVis Group at the University of British Columbia and the Bedford lab at the Fred Hutch Institute. The authors would also like to acknowledge their funding sources. AC is supported by a CIHR Vanier Scholarship, JG by the Canada Research Chairs Program and a Michael Smith Foundation for Health Research Scholar award, and TM by the NSERC Discovery Program.

## Author Contributions

AC, JG, and TM devised and interpreted the analysis and jointly wrote the paper.

